# Spatiotemporal changes in secondary metabolites during graft formation in grapevine reveal tissue-specific accumulation of metabolites in necrotic and callus tissues

**DOI:** 10.1101/2023.03.09.531860

**Authors:** Grégoire Loupit, Josep Valls Fonayet, Marcus Daniel Brandbjerg Bohn Lorensen, Céline Franc, Gilles De Revel, Christian Janfelt, Sarah Jane Cookson

## Abstract

Grafting is widely used in horticulture, shortly after grafting, callus tissues appear at the graft interface and the vascular tissues of the scion and rootstock connect. The graft interface contains a complex mix of tissues, we hypothesized that each tissue have is own metabolic response to wounding/grafting and accumulate different metabolites at different rates. We made intact and wounded cuttings and grafts of grapevine, and then measured changes in bulk secondary metabolite concentration and used metabolite imaging to study tissue specific responses. We show that some metabolites rapidly accumulate in specific tissues after grafting, e.g. stilbenes accumulate in necrotic tissues surrounding mature xylem vessels and gradually oligomerize over time. We also observe that some metabolites accumulate in the newly formed callus tissue at the graft interface and identify genotype-specific responses. Here we reveal the spatiotemporal dynamics of metabolite changes occurring during graft union formation for the first time. The rapid accumulation of stilbenes in the tissues damaged during the grafting process could be a plant defence mechanism, as stilbenes have antioxidant and anti-fungal capacities. The increasing oligomerization of stilbenes often occurs in response to plant stresses (via unknown mechanisms), but it potentially increases antioxidant activity.

**Brief summary:** Secondary metabolites accumulate after wounding and grafting in plants yet we have limited knowledge of tissue specific accumulation patterns and temporal dynamics. We show that stilbenes accumulate specifically in necrotic tissues and oligomerize over the time, whereas other compounds accumulate in the newly formed callus tissues. This suggests that these compounds have different roles in wounding healing and grafting.

## Introduction

Grafting is a technique widely used in horticulture, particularly for perennials, and is increasingly used for annual plant production (Rouphael, Schwarz, Krumbein, & Colla, 2010). This ancestral technique makes it possible, by the combination of two tissues from two different plants, to form a single plant with the agronomical advantages of the scion and the rootstock genotypes (Mudge, Janick, Scofield, & Goldschmidt, 2009). The advantages of using grafting in agriculture are that we can independently select for desirable root and shoot traits.

Grafting uses the intrinsic capacities of plants to heal, which includes cell proliferation and dedifferentiation, allowing tissue reconnection, which is particularly at the vascular level (Hartmann, Kester, Davies Jr, & Geneve, 2011; Melnyk, 2017). Grafting starts with a mechanical injury, which triggers changes in plant metabolism and signalling, such as the production of reactive oxygen species and the activation of defence and wound healing mechanisms. In general, secondary metabolites accumulate at the graft interface during the months and years after grafting (Assunção, Pinheiro, et al., 2019; Canas, Assunção, Brazão, Zanol, & Eiras-Dias, 2015; Musacchi, Pagliuca, Maddalena, Piretti, & Sansavini, 2000; Usenik, Krška, Vičan, & Štampar, 2006) and presumably these molecules have a role in plant defence mechanisms (Chong, Poutaraud, & Hugueney, 2009; L. Yang et al., 2018). In addition to biochemical changes, callus cells often proliferate at the graft union, symplastic connections are made via plasmodesmata and the phloem and xylem connect between the scion and rootstock (Hartmann et al., 2011).

Although widely used, grafting is sometimes unsuccessful. These grafts are characterized by a low survival rate, weak graft unions and poor growth. Poor grafting success can have multifactorial origins such as the combination of certain scion/rootstock genotypes, poor grafting technique, the presence of pathogens or due to climatic conditions (Hartmann et al., 2011). Depending on the scion/rootstock combinations, the grafting success can therefore vary highly as well as from year to year. The genetic distance between the two grafted plants seems to be a determining factor in grafting success, however, tobacco (*Nicotiana benthamiana*) is high graft compatible with many different species, even with very distantly related ones (Notaguchi et al., 2020), although whether these grafts are truly functional, with connected vascular tissues, has not been determined. The origins of graft incompatibilities are difficult to elucidate, but research in this direction is essential because the use of new rootstocks could be a key factor of adaptation climate change for many agroecosystems (Ollat et al., 2015).

Many studies have attempted to identify metabolite markers of a successful graft union formation or graft incompatibility (Assunção, Pinheiro, et al., 2019; Canas et al., 2015; DeCooman, Everaert, Curir, & Dolci, 1996; Errea, 1998; Hudina, Orazem, Jakopic, & Stampar, 2014; Loupit et al., 2022), but with limited success (Loupit & Cookson, 2020). To our knowledge, no study has examined the changes in secondary metabolite concentration occurring around graft interface in the hours and days after grafting in any plant species, which is one of the objectives of the current study. In addition, the spatial distribution of secondary metabolites in the different woody tissues has been little studied in any species, although a small number of papers have been published in recent years. For example, laser micro-dissection, dissection and metabolite imaging has been used to characterise the metabolite profiles of different tissues in tree species (Abreu et al., 2020; Hu et al., 2022; G. Yang, Liang, Zhou, Wang, & Huang, 2020), but few have examined metabolites in the bark and pith/heartwood. Furthermore, we know little of the tissue-specific metabolite changes triggered by wounding or grafting woody tissues in any plant species.

We have previously shown that certain metabolites, particularly stilbenes, accumulate at the graft interface of grapevine at least one month after grafting (Loupit et al., 2022; Prodhomme et al., 2019), but the spatiotemporal dynamics underlying these metabolite changes are not known. In this study, we combine a time course experiment on bulk tissue (using High Performance Liquid Chromatography coupled with a Triple Quadrupole Mass Spectrometer (HPLC-QqQ)) and metabolite imaging (using matrix-assisted laser desorption/ionization-mass spectroscopy / MALDI-MS) to study the tissue specific changes in metabolites occurring during graft union formation.

To characterise the temporal changes in secondary metabolites during grafting, we quantified several secondary metabolites above, below (representing the scion and the rootstock in grafted combinations) and at the graft interface at 0, 4 hours, 1, 3, 6 and 15 days after grafting (DAG) in two hetero-grafts (*Vitis vinifera* cv. Pinot Noir (PN) on *Vitis riparia Michaux* cv. Riparia Gloire de Montpellier (RGM) and PN on *V. berlandieri* x *V. rupestris* cv. 140 Ruggeri (140Ru)). These two rootstocks were chosen for their differences in graft development: the rootstock 140Ru often produces grafts with a weak graft junction and large callus, whereas RGM is highly graft compatible (Pl@ntGrape Database). In addition, to understand the genotype-specific metabolite responses to grafting, we studied the three homo-grafted controls of these genotypes (PN/PN, RGM/RGM and 140Ru/140Ru), as well as wounded and intact PN cuttings. We measured 41 secondary metabolites of interest (20 stilbenes, 14 flavanols, caftaric acid, coutaric acid, quercetin-3-glucoside, quercetin-3-glucuronide, naringenin, naringenin glucoside and taxifolin) in these samples.

The graft interface is complex, mixing two different genotypes, composed of different tissues (xylem, phloem, bark and pith) which are both intact and damaged, as well as the newly formed callus. Because of this, we hypothesized that each tissue have is own metabolic response to grafting and accumulate different metabolites. Therefore, to understand in which tissues the metabolites identified the time course experiment accumulate, we used Matrix Assisted Laser Desorption Ionization mass spectrometry imaging (MALDI-MSI) to visualize metabolite spatial distribution at the graft interface at 16 and 30 DAG. MALDI-MSI is a relatively recent tool and is being increasingly used to study the distribution of metabolites in plant tissues (Bjarnholt, Li, D’Alvise, & Janfelt, 2014). So far few studies have been done on either woody tissues (Hu et al., 2022) or grapevine (Becker, Carré, Poutaraud, Merdinoglu, & Chaimbault, 2014).

## Materials and methods

### 1. Plant material

Overwintering grapevine canes were used to make the grafts and cuttings (Table 1); PN and 140 Ru were supplied by the Chambre de l’Agriculture de l’Aude (Carcassonne, France), and RGM was supplied by Mercier (Vix, France). The HPLC-QqQ analysis was done in 2020 and the MALDI-MSI was done in 2022. Grapevine canes were stored in a fridge in 1 m long sections until shortly before grafting, then one bud scions and de-budded rootstocks of about 30 cm in length were prepared. One day before grafting, the scions and rootstocks were rehydrated by soaked in water at room temperature for 4 hours.

**Table 1:**
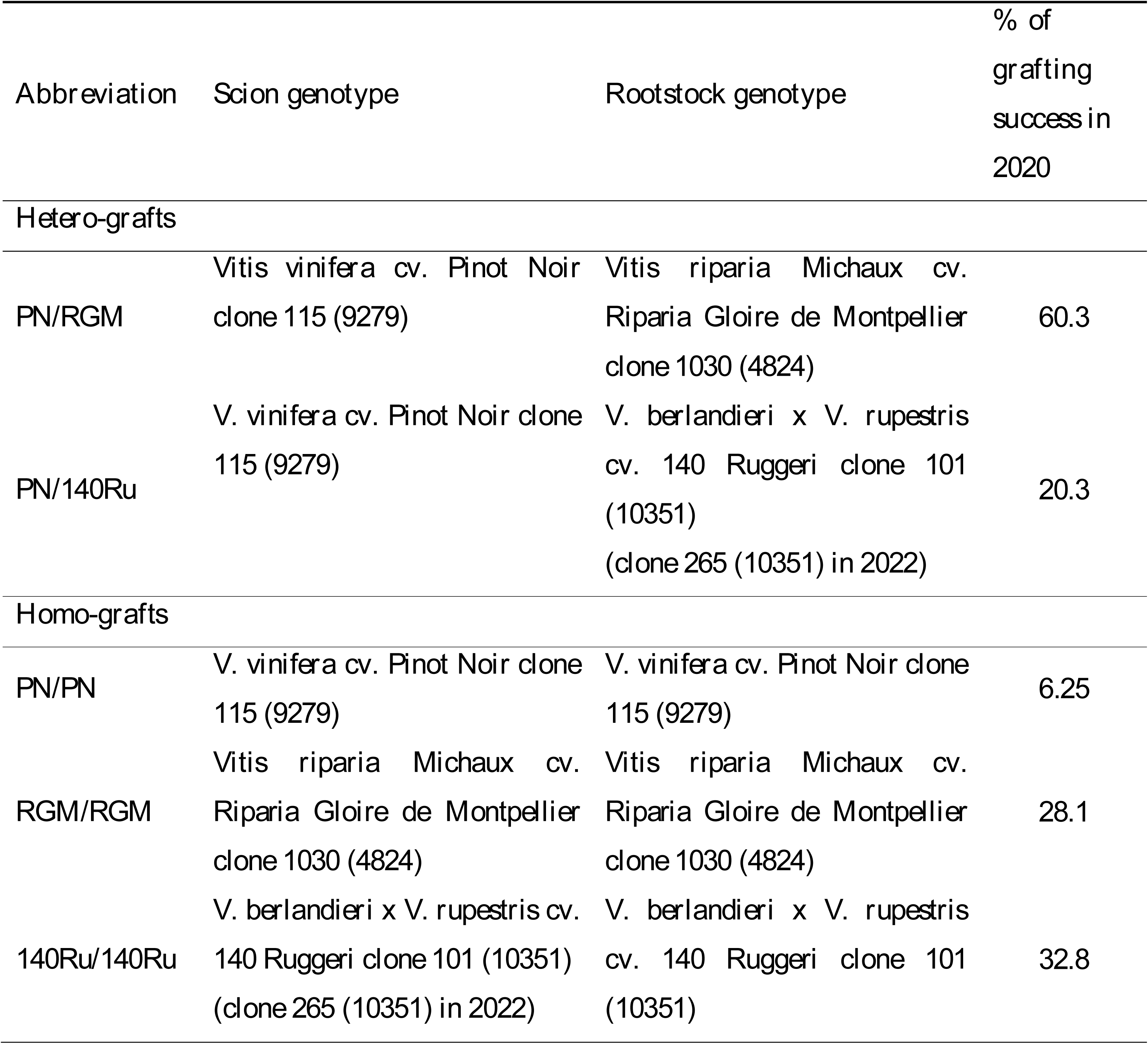
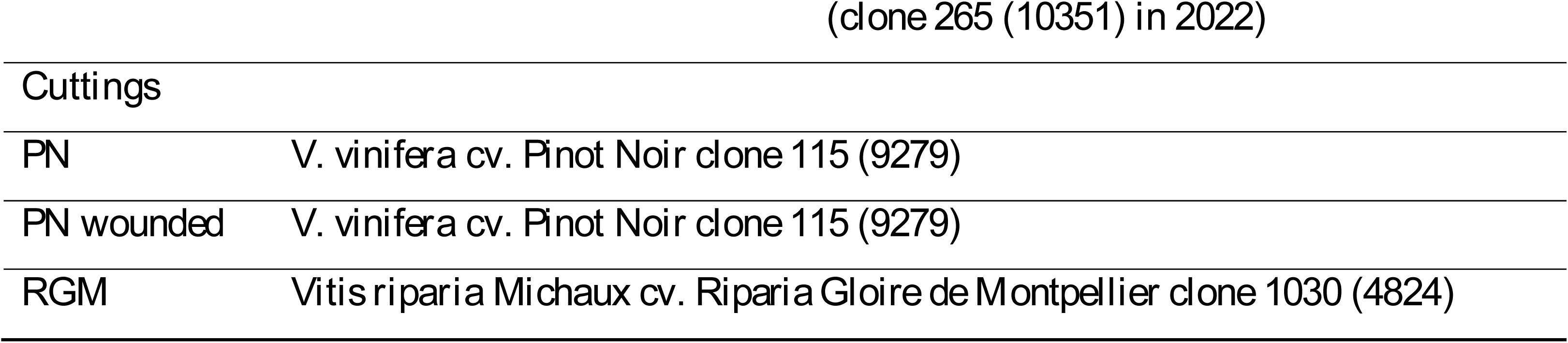
The grafted combinations used in both experiments, grafting success in 2020 and Vitis International Variety Catalogue numbers in brackets.

### 2. Grafting procedure

For the time course experiment, 250 grafts were made for the five scion/rootstock combinations and 100 cuttings were made for each treatment. For the MALDI-MSI experiment, 100 grafts were made for each scion/rootstock combinations and 60 cuttings were made of RGM. All grafts were made using an Omega blade (Omega Star, Chauvin, France). Then, all plants were dipped in melted wax (Staehler Rebwachs pro with 0.0035% of dichlorobenzoic acid, Chauvin) and put in plastic boxes for 5 d at room temperature and then placed in callusing room (28 °C and high humidity) with 2 cm of water in crates for 20 to 30 d depending on scion/rootstock combination.

For the time course experiment, at each date (0 and 4 h after grafting, and 1, 3, 6 and 15 DAG), five pools of five grafts and three pools of four cuttings were harvested randomly. Tissues were sampled at the graft interface, and 1 cm above and below the graft interface (without nodes) for the grafts, and 1 cm above, 1 cm below and at wounded interface for cuttings (in the case of PN intact cuttings, three different parts were sampled to correspond to the position of above, below and at the graft interface) (Figure 1). Grafts remaining after harvesting were dipped in melted wax for a second time and planted in pots with fertigation (N=34.3; P=17.7; K=140.4; Ca=74.9; Mg=16.8; S=41.4; Cl=90.1 mg L^-1^). On the 2^nd^ of January 2021, the mechanical resistance of the graft union was tested to calculate grafting success rates for the five scion/rootstock combinations. Grafts were deemed successful if they did not break during the mechanical resistance test (Table 1).

**Figure 1:**
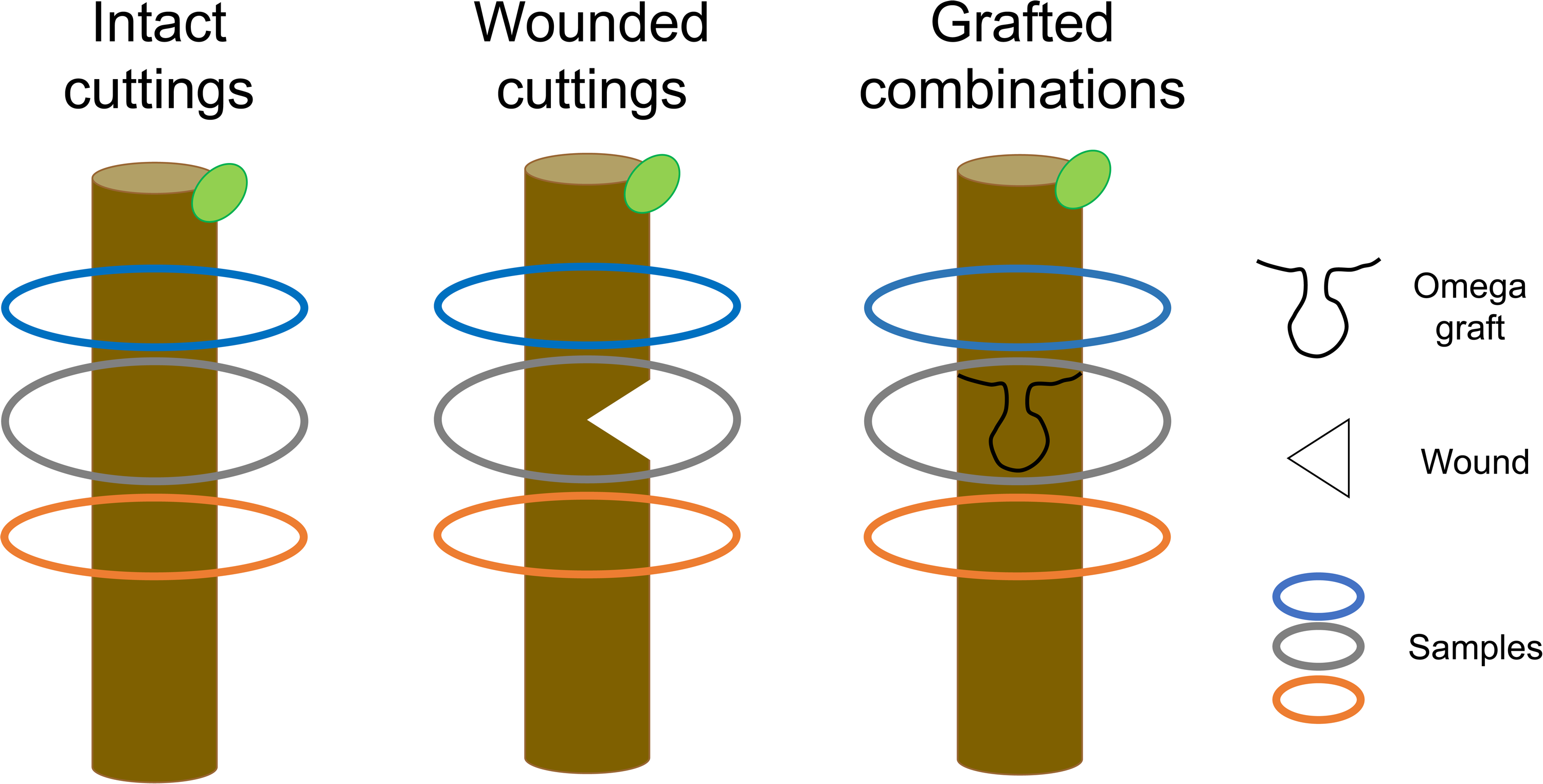
Experimental design. Sampling were taken from intact and wounded cuttings, and grafted plants 0 and 4 h, and 1, 3, 6 and 15 d after grafting (DAG). At each time point, five pools of five grafts and three pools of four cuttings were harvested randomly.

For the MALDI-MSI experiment, two RGM cuttings and two grafts of each hetero-graft and homo-graft studied at 0, 16 and 30 DAG. All samples were immediately snap frozen in liquid nitrogen and kept at −80 °C until analysis.

### 3. Chemicals and standards

All standards used came from Extrasynthesis (France) except for some stilbene monomers (*trans-*astringin and *trans*-isorhapontin), stilbene dimers (*trans-*ε-viniferin, *trans*-ω-viniferin , *trans*-δ-viniferin, pallidol, parthenocissin A, vitisinol C and ampelopsin A), stilbene trimers (miyabenol C and α-viniferin), and stilbene tetramers (hopeaphenol, isohopeaphenol, r2-viniferin and r-viniferin) which have been purified in the MIB laboratory. 1,5-Diaminonaphthalene (DAN, MALDI matrix) and carboxymethylcellulose sodium salt (CMC, embedding material), where both purchased from Sigma-Aldrich (Copenhagen, Denmark).

### 4. Analysis by UHPLC-QqQ

Before extraction, samples were ground to powder in a ball mill (MM400 RETSCH) at 30 Hz during 30 s. The extraction protocol was the same as used by Loupit *et al*. (2022) (250 mg of ground powder extracted with 4 mL of methanol during 15 mins in an ultrasound bath) (Loupit et al., 2022). Then, polyphenols analysis was done as described by Loupit *et al*. (2020) except that some additional compounds were added (Table S1) an alternative column was used (Agilent ZORBAX RRHD SB-C18 (2.1 mm x 100 mm, 1.8 μm)) (Loupit et al., 2020). Standards were used to quantify the different compounds (concentration ranged from 0.01 to 20 mg L^−1^, to 50 mg L^−1^ for catechin, epicatechin, B1, hopeaphenol, r-viniferin and to 200 mg L^−1^ for *trans-*ε-viniferin) except for procyanidin B3, B4, *cis*-piceid, *cis*-astringin and coutaric acid were quantified as procyanidin B1, B2, *trans*-piceid, *trans*-astringin and caftaric acid respectively. Furthermore, one flavanol dimer and two flavanol trimers were given the names dimer, trimer2 and trimer3 respectively, and quantified as procyanidin B2 (for the dimer) and procyanidin C1 (for the trimers). All concentration found are given as g kg^−1^ FW (Data S1).

### 5. Analysis by MALDI-MSI

Graft interfaces, cuttings and wood samples, were embedded in a 2% carboxy methyl cellulose (CMC) solution with Milli-Q water and kept at −80 °C. Cryosectioning was performed as described by Granborg *et al*. (2022) by using a Leica CM3050S cryo-microtome (Leica Microsystems, Wetzlar, Germany) at −25°C to make 20 μm thick sections adapted from Kawamoto *et al*. (2003), harvested at the middle of the wood (Granborg, Kaasgaard, & Janfelt, 2022; Kawamoto, 2003). Sections were attached to microscope slides using double-sided graphite tape (Electron Microscopy Sciences, Hatfield, PA, USA) and then stored at −80 °C until analysis.

Before matrix application, sections were dried for 20 min in a vacuum desiccator. Then, 300 μL of matrix solution (3.3 mg mL^−1^ of 1,5-diaminonaphthalene (DAN) in 90:10 methanol:H O (v/v)) was applied at 30 μL min using a pneumatic sprayer (the distance between sprayer and sample was 10 cm with a gas pressure of 2.0 bar and the sample was rotated at 600 rpm).

MALDI-MSI analysis was performed on a Thermo QExactive Orbitrap mass spectrometer (Thermo Scientific GmbH, Bremen, Germany) equipped with an AP-SMALDI10 ion source (TransMIT GmbH, Giessen, Germany) operated at a mass resolving power of 140 000 at *m/z* 200, with a lock-mass from matrix peak of DAN (*m/z* 311.1302). The analysis was performed in negative-ion mode with a pixel size of 35 μm and using a range analysis between 100 to 1000 *m/z*. Finally, RAW files were converted to imzML files with RAW2IMZML software (TransMIT GmbH, Giessen, Germany) and images were made with MSiReader ver. 1.02 (Bokhart, Nazari, Garrard, & Muddiman, 2018; Robichaud, Garrard, Barry, & Muddiman, 2013) with normalization to the total ion current (TIC) and a mass tolerance of 5 ppm.

### 6. Statistics analysis

HeatMaps were made on software R (version 4.0.4) and RStudio (version 1.2.5019) using gplots package. Statistical tests (T-test) were done online on BioStatFlow v.2.9.5 © INRAE 2022 to identify metabolites that accumulates specifically at the graft interface. Stars, positioned on time course data and HeatMaps, indicates only a significant difference between scion and graft interface, and between graft interface and rootstock. Significance threshold set at *p*-value < 0.05. All *p*-value for each grafted combinations and cuttings are given in Data S2.

## Results

### Stilbenes accumulate at the graft interface

To investigate the temporal changes in secondary metabolites occurring during graft union formation in grapevine, we measured 41 metabolites in two hetero-and three homo-grafted scion/rootstock combinations above, below and at the graft interface during the first 15 DAG. To identify metabolite profiles specific to grafting, we also compared the secondary metabolite responses to grafting and wounding in the PN genotype. In addition, we measured metabolites in PN cuttings at the same location as the tissues harvested for above, below and at the graft interface as intact controls. The sum of the flavanols and stilbenes in the different tissues provides an overview of the metabolomic changes occurring (Figure 2). Total flavanol and stilbene concentration did not change in the stems of intact PN cuttings over the 15 d period. However, in all grafting or wounding treatments, stilbenes accumulated at the graft interface/wound site, while the quantity of flavanols remained stable or slightly decreased over time (Figure 2). Total stilbene concentration reached a maximum at 6 DAG at the graft interface/wound site in all treatments. The three genotypes studied had intrinsically different concentrations of stilbenes at T0, with the highest concentrations in RGM and the lowest in 140Ru. In addition, the rate of stilbene accumulation at the graft interface was different between the three homo-grafts: the stilbene accumulation at the graft interface was already significant at 1 DAG for 140Ru/140Ru and RGM/RGM, whereas it was significant only at 3 DAG for PN/PN. Furthermore, the relative increase at the graft interface in relation to the surrounding woody tissues was higher in 140Ru/140Ru (approximately two-fold) compared to the other two homo-grafts (approximately 1.3-fold). In addition, total stilbene content below and above the graft interface remained stable during the 15 DAG in 140Ru/140Ru, but increased by 1.7 and 1.5-fold in PN/PN and RGM/RGM homo-grafts respectively. The maximum total stilbene accumulation at the graft interface was higher in PN/PN homo-grafts compared to wounded PN cuttings, while the amount found at 1 DAG was similar in both conditions.

**Figure 2:**
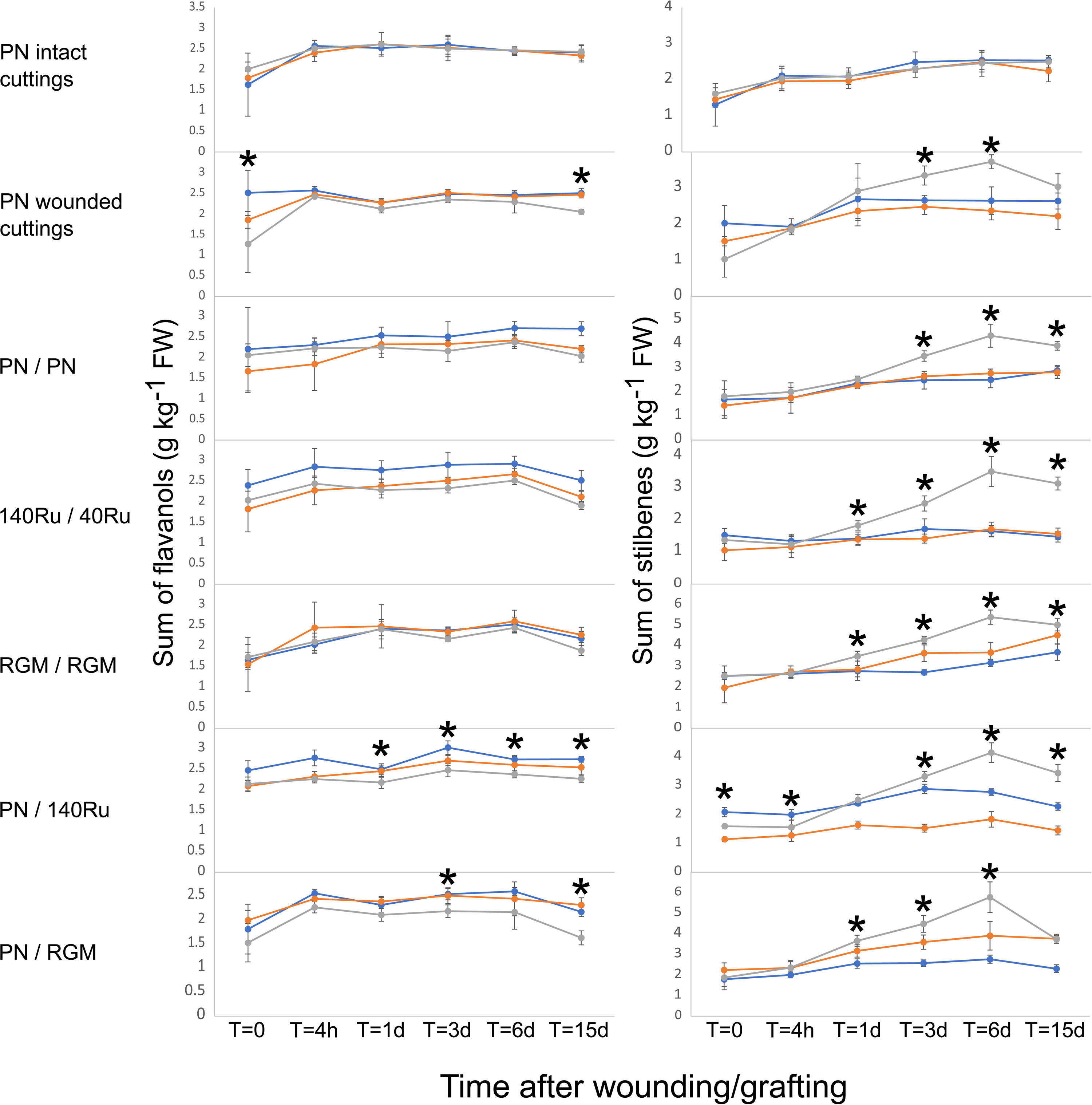
Effect of wounding and grafting on the sum of flavanols and stilbenes at 0 and 4 h, and 1, 3, 6 and 15 d after grafting/wounding grapevine stems. Intact and wounded cuttings of PN (*V. vinifera* cv. Pinot Noir) were studied, as well as various homo-and hetero-grafts of three grapevine genotypes (PN, RGM (*V. riparia* Michaux cv. Riparia Gloire de Montpellier) and 140 Ru (*V. berlandieri x V. rupestris* cv. 140 Ruggeri). Samples were taken above (blue), below (orange) and at the graft interface (grey). Symbols represent means ± the standard deviation (n=5 for grafts and n=3 for cuttings). Stars indicate a significant difference between scion and graft interface, and between graft interface and rootstock after a T-test analysis with a significance threshold set at *p*-value < 0.05.

The concentration of total stilbenes in the hetero-grafts depended on both the genotype-specific differences in stilbene concentration and the accumulation of total stilbenes at the graft interface and showed similar patterns to the homo-grafts.

The accumulation of the different stilbene compounds at the graft interface of the two hetero-grafts was globally similar except for certain compounds such as isorhapontigenin, pallidol, α-viniferin or miyabenol C (Data S1). These differences seemed to be mainly due to genotype-specific differences in metabolite responses to wounding/grafting. For example, isorhapontigenin and α-viniferin are more highly accumulated at the graft interface of PN/140 than PN/RGM, this appears to be primarily due to the high levels of accumulation of these compounds by the 140Ru genotype. Similarly, pallidol and miyabenol C are more highly accumulated at the graft interface of PN/RGM than PN/140Ru, this appears to be related to their high accumulation at the graft interface of RGM/RGM (Data S1).

### Stilbenes accumulate from 1 DAG and oligomerize over the time

To gain further insights into the accumulation of stilbenes at the graft interface, the sum of the different monomers (7), dimers (7), trimers (2) and tetramers (4) was calculated (Figure 3). Concerning the intact PN cuttings, the concentration of the different stilbenes did not show significant differences in the sites of “above”, “below” and at the “graft interface”. However, the concentrations of stilbene monomers and dimers increased from 1 DAG to 6 DAG, trimers from 6 DAG and tetramers increased at 15 DAG (Figure 3). This shows that stilbene tissue concentrations increased in concentration upon transfer from the cold room to warm, growing conditions in the wood of PN, but variation concentrations were globally negligible compared to wounded and grafted conditions.

**Figure 3:**
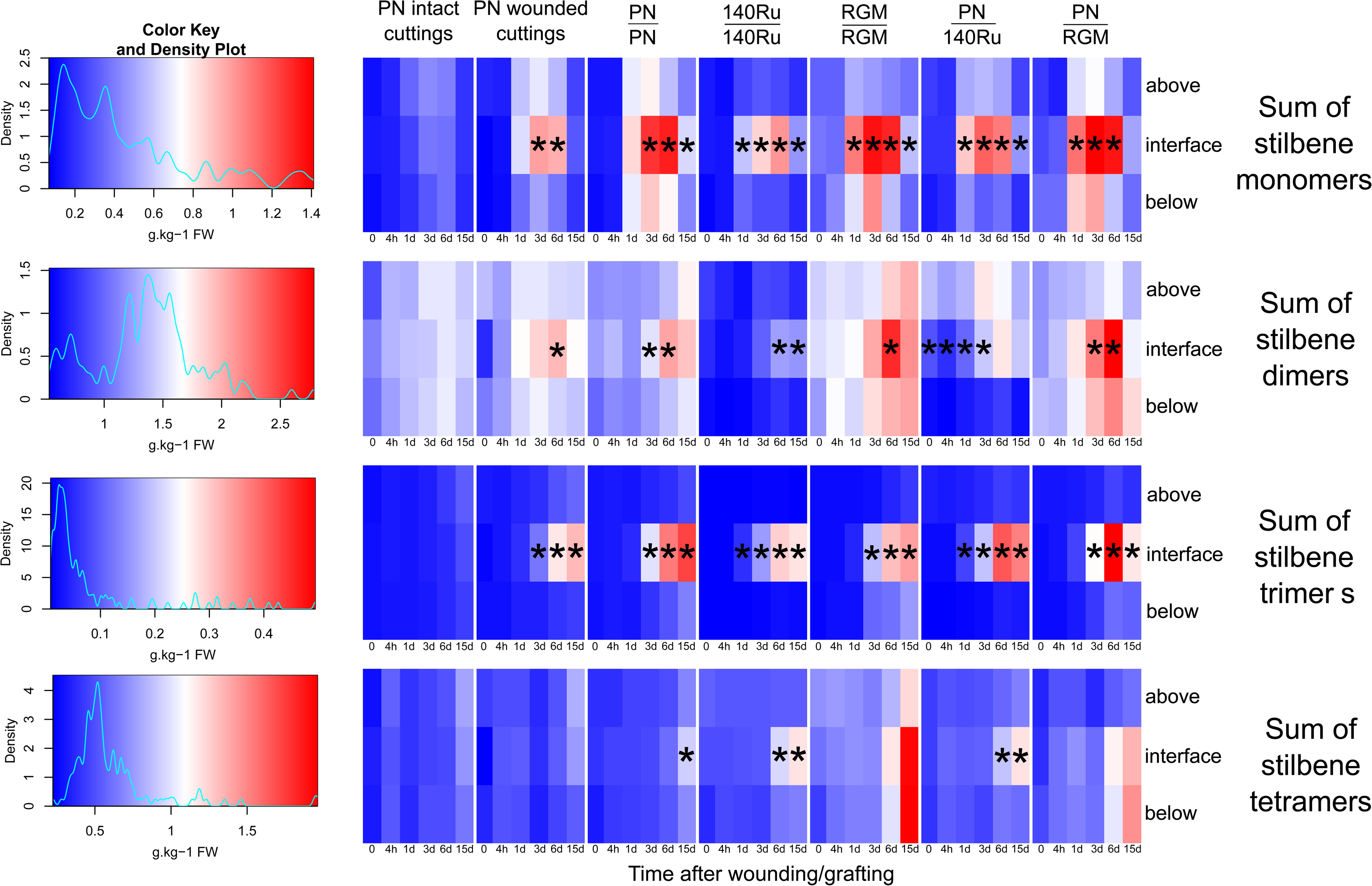
Heatmaps of the effect of wounding and grafting on sum of stilbene monomers, dimers, trimers and tetramers at 0 and 4 h, and 1, 3, 6 and 15 d after grafting/wounding grapevine stems. Cuttings and grafts as described in legend for Fig. 1. Stars indicate a significant difference between scion and graft interface, and between graft interface and rootstock after a T-test analysis with a significance threshold set at *p*-value < 0.05.

Stilbene monomers, dimers and trimers accumulated to much higher levels at the wound site and graft interface in all scion/rootstock combinations. The accumulation of monomers was continuous and significant from 1 DAG in the 140Ru and RGM homo-grafts and in the two hetero-grafts, and from 3 DAG in wounded cuttings and PN homo-grafts. This accumulation pattern was largely due to the accumulation of resveratrol, the major stilbene monomer in these tissues (Data S1). However, at 15 DAG, the quantity of monomers decreased significantly in comparison to 6 DAG; this decrease may be due to the metabolization of these compounds into more complex metabolites such as oligomeric forms. In general, the accumulation of dimers, trimers and tetramers at the graft interface occurred later in the time course than the accumulation of monomers (Figure 3). Tetramers only accumulated at the graft interface towards the end of the time course and only in the PN/PN, PN/140 Ru and 140Ru/140Ru combinations (Figure 3).

Stilbene dimers were present in large quantities in all the tissues studied, due to the high content of *trans*-ε-viniferin (Data S1). The rate of accumulation of *trans*-ε-viniferin at the graft interface was strongly dependent on the scion and rootstock genotype. This is because stilbene dimer concentration was much lower in wood of 140Ru than the other two genotypes (Figure 3). Furthermore, the relative increase in dimer concentration at the graft interface depended on the scion/rootstock genotypes; for example, in 140Ru homo-grafts, dimer concentration doubled from 0 to 15 DAG, whereas in RGM homo-grafts maximum dimer concentration was at 6 DAG and the relative increase was approximately 40 % from 0 to 6 DAG (Data S1).

The accumulation of trimers at the interface began at 3 DAG (except for combinations with the 140Ru genotype where it began from 1 DAG) and increased until 15 DAG (at least by 2-fold) except for PN/RGM (Figure 3). Finally, only 140Ru/140Ru and PN/PN homo-grafts and PN/140Ru hetero-grafts showed an increase of stilbene tetramers at 6 or 15 DAG at the graft interface compared to above and below it. Although tetramers did not accumulate at the graft interface of RGM/RGM homo-grafts and PN/RGM, they accumulated below the graft interface (Figure 3), especially for hopeaphenol and isohopeaphenol (Data S1).

### Naringenin, taxifolin and compounds with a gallate residue also accumulate at the graft interface

The quantification of phenolic acids, flavonols and flavanols during the first 15 DAG showed significant differences between the genotypes and/or graft combinations studied, but no specific accumulation at the graft interface or wound site occurred except for naringenin and its glucoside form, taxifolin, and flavanol compounds with a gallate residue (Figure 4, Data S1). These compounds were at the same concentrations in the different parts of intact PN cuttings, showing that the accumulation was specific to wounding or grafting despite the very low quantities measured. In all grafts and wounded cuttings, a strong accumulation of naringenin at the wound site began from 1 DAG (for example in 140Ru/140Ru homo-grafts at 1 DAG its concentration was 3 times higher at the interface compared to the scion and rootstock). However, this strong and rapid accumulation decreased over time until it was no longer significant at 15 DAG in wounded PN cuttings, PN/PN and RGM/RGM homo-grafts, as well as in PN/RGM hetero-grafts (Figure 4). The accumulation pattern was similar for taxifolin except for 140Ru/140Ru homo-grafts, which accumulate high concentrations of taxifolin that remain high for longer than the other scion/rootstock combinations. Flavanols with a gallate residue also accumulated at the graft interface and wounding site, but only from 3, 6 or 15 DAG (Figure 4), the differences observed were largely due to differences in the accumulation of epigallocatechin gallate (Data S1).

**Figure 4:**
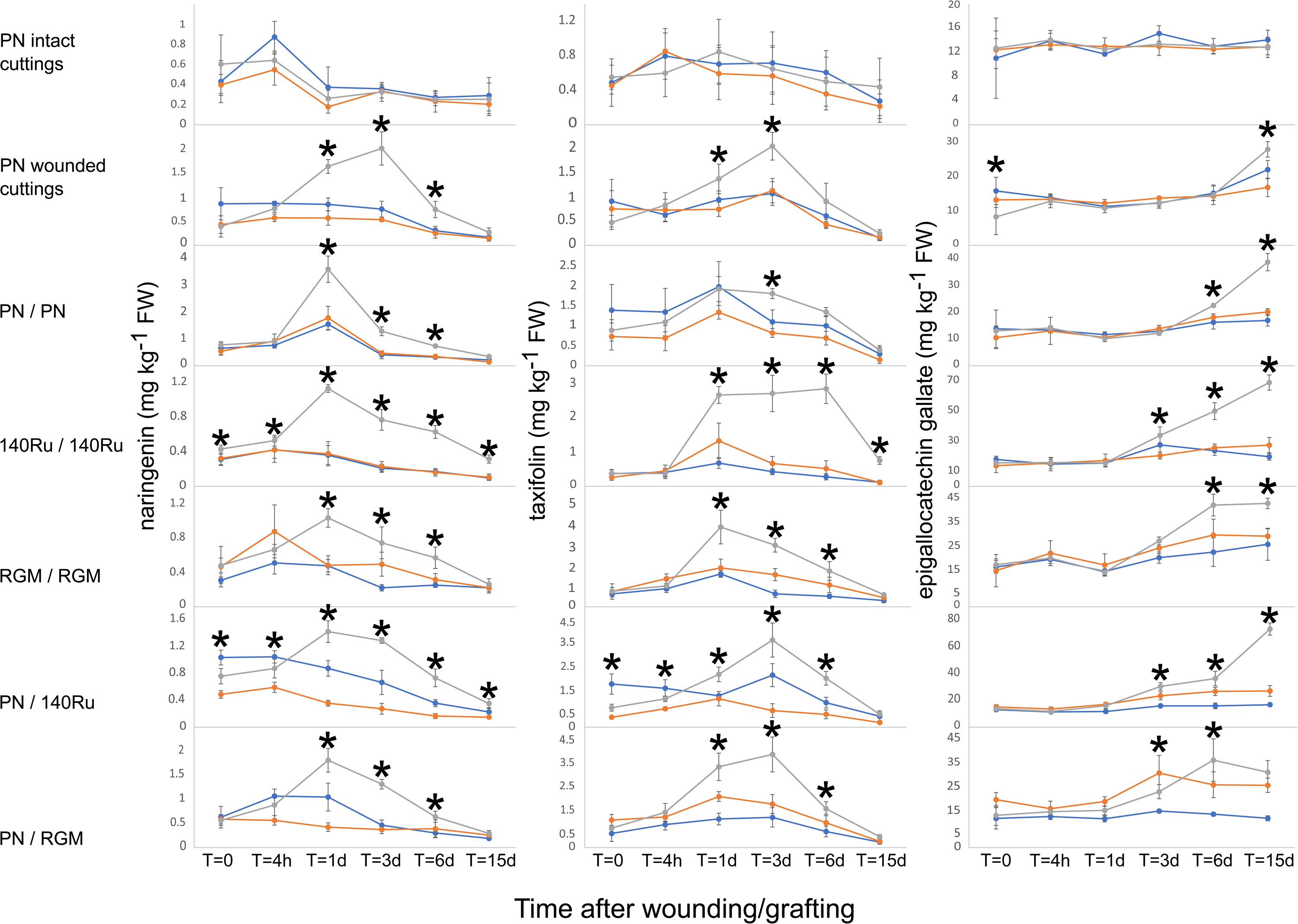
Effect of wounding and grafting on naringenin, taxifolin and epigallocatechin gallate concentration at 0 and 4 h, and 1, 3, 6 and 15 d after grafting/wounding grapevine stems. Cuttings and grafts as described in legend for Fig. 1. Symbols represent means ± the standard deviation (n=5 for graft combination and n=3 for cuttings). Samples were taken above (blue), below (orange) and at the graft interface (grey). Stars indicate a significant difference between above and at the graft interface, and between below and at the graft interface (T-test analysis, *p*-value < 0.05).

### The spatial distribution of secondary metabolites in different tissues of wood stems of grapevine

The use of MALDI-MSI allowed us to observe the spatial distribution of different compounds at different times after grafting (at 0, 16 and 30 DAG). In PN, 140Ru and RGM wood (at 0 DAG), stilbenes (such as resveratrol, dimers, α-viniferin and tetramers) were found between the xylem part and the pith, as well as between the bark and phloem. In general, the distribution of these metabolites between the different tissues was similar between the different genotypes. However, in PN (Tables S4 and S5), resveratrol, stilbene dimers, α-viniferin and tetramers are more highly concentrated in the pith, and tetramers seemed to be at lower concentration in the bark compared to 140Ru (Tables S2 and S3) and RGM (Tables S6 and S7). Concerning flavanol monomers (catechin, epicatechin) and flavanol dimers (B1, B2, B3, B4, etc…), we found a similar tissue specific distribution to that of the stilbenes, except that flavanols were more highly concentrated in the pith and in their distribution was more homogeneous. However, for PN, flavanols were also present in high concentrations in the xylem and phloem tissues (Tables S4 and S5). The distribution of taxifolin and naringenin was similar to that of the other compounds studied, with high concentrations in the pith and bark, and high concentrations of naringenin also in the xylem area of PN (Table S4 and S5). Despite these small differences in metabolites distribution patterns, overall the localisation of the different metabolites of interest was similar between genotypes.

### Stilbenes accumulate in wounded tissues while naringenin, taxifolin and epigallocatechin gallate accumulate in callus

In agreement with the time course data, flavanol monomers and dimers did not accumulate at the graft interface in any of the grafts studied. These compounds were distributed in the same fashion as in the cuttings at 0 DAG. However, these compounds accumulate to low levels in the newly formed callus at 30 DAG (Table S2-11).

Concerning resveratrol, we observed an accumulation at 16 DAG along the zone of damaged tissue at the graft interface, which corresponds to the omega shape of the grafting machine (Figure 5a and 6a). This accumulation was clearly visible, particularly in the xylem parenchyma tissues in the PN and RGM genotypes, and in hetero-graft and homo-graft combinations. Conversely, in 140Ru genotype, the accumulation was throughout the xylem zone, but also in the pith (Figure 5a and Tables S2 and S3). The localised accumulation of resveratrol in the damaged tissue of the graft interface was generally less visible at 30 DAG in most grafts, except for RGM/RGM homo-grafts (Table S6 and S7). This agrees with time course data in which the accumulation of resveratrol decreases from 6 DAG. Stilbene dimers had similar distribution to resveratrol in hetero-graft combinations (Tables S8-11) but seemed to accumulate less specifically at the interface in homo-grafted controls (Tables S2-7) whatever the time after grafting studied.

**Figure 5:**
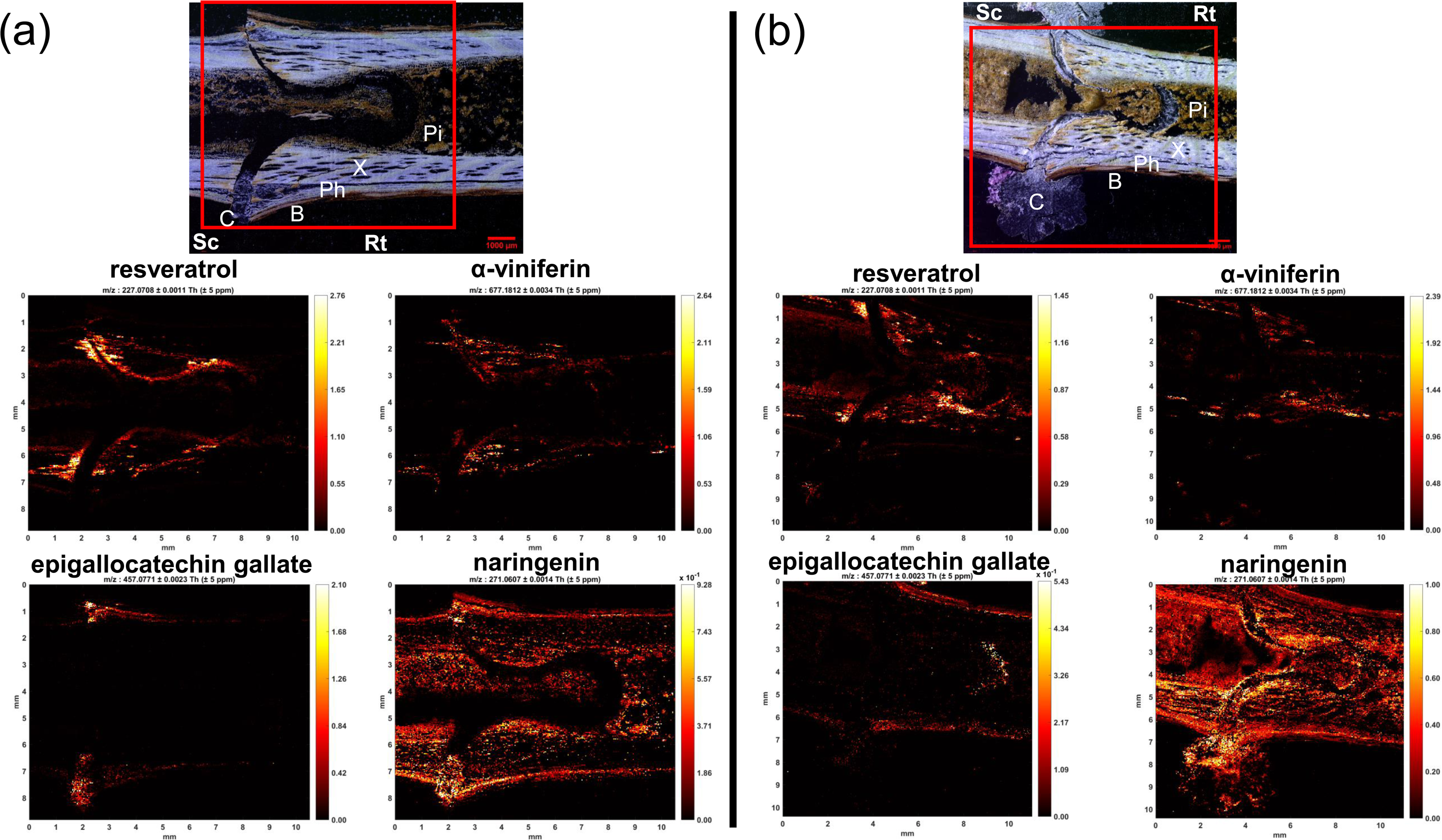
Photographs of *V. vinifera* cv. Pinot Noir/*V. berlandieri x V. rupestris* cv. 140 Ruggeri hetero-grafts sections at (a) 16 and (b) 30 DAG and their MS images generated for *m/z* 227.0708, *m/z* 677.1812, *m/z* 457.0771 and *m/z* 271.0607 corresponding to resveratrol, α-viniferin, epigallocatechin gallate and naringenin respectively. The red boxes correspond to the area analysed. Sc, scion; Rt, rootstock; C, callus; B, bark; Ph, phloem; X, xylem; Pi, pith.

**Figure 6:**
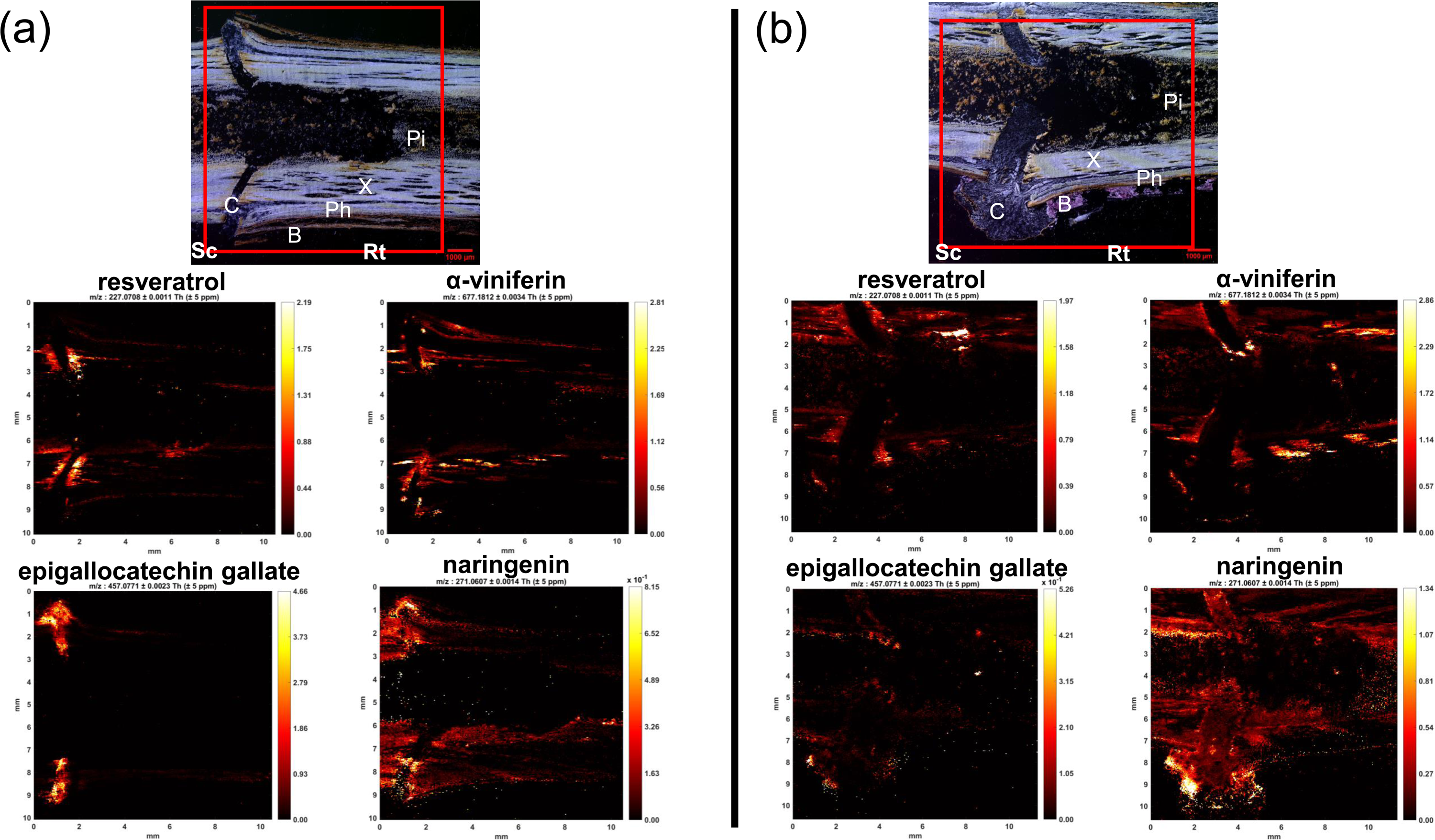
Photographs of *V. vinifera* cv. Pinot Noir/*V. riparia* Michaux cv. Riparia Gloire de Montpellier hetero-grafts sections at (a) 16 and (b) 30 DAG and their MS images generated for *m/z* 227.0708, *m/z* 677.1812, *m/z* 457.0771 and *m/z* 271.0607 corresponding to resveratrol, α-viniferin, epigallocatechin gallate and naringenin respectively. The red boxes correspond to the area analysed. Sc, scion; Rt, rootstock; C, callus; B, bark; Ph, phloem; X, xylem; Pi, pith.

Concerning the concentration of stilbene trimers such as α-viniferin, in grafted tissues α-viniferin was most concentrated in the xylem part, along the damaged tissues as well as around the xylem vessels, but it was absent from the pith (Figure 5 and 6). In 140Ru genotype, α-viniferin was highly concentrated between the xylem part and the pith, whereas in PN and RGM genotypes, this observation was not so apparent. Concerning the stilbene tetramers, their distribution is similar to those of other stilbenes: an accumulation was visualized along the interface at 16 DAG in the hetero-grafts, but it seemed to disappear at 30 DAG. Moreover, unlike dimers or resveratrol and the wood at 0 DAG, the presence of stilbene tetramers was not visible between the bark and the phloem part (Tables S8-11). In some cases, especially in PN genotype, resveratrol, dimers and trimers also accumulated at the outer edge of the callus.

The visualisation by MALDI-MSI showed that epigallocatechin gallate, naringenin, and taxifolin accumulate at the graft interface at 16 DAG (Figure 4, Tables S2-11). In all grafts, epigallocatechin gallate was found specifically accumulated in callus at 16 DAG, however at 30 DAG it has a more diffuse distribution across the tissues (Table S2-11). Naringenin or taxifolin were easily visible with MALDI-MSI and were widely distributed in the tissues, these compounds accumulated to high levels in the callus tissues (Tables S2-11), but also along the tissues damaged during the grafting process, in particular at 30 DAG in the pith in PN/140Ru hetero-grafts and RGM homo-grafts (Figure 5b, Table S6 and S7). Presumably the concentrations of naringenin and taxifolin at 16 DAG were much lower than those at 1 DAG when these compounds are at their highest concentration (Figure 4).

## Discussion

The objective of this study was to characterise the spatiotemporal changes in secondary metabolites occurring at the graft interface during graft union formation in grapevine, a woody perennial crop species. This is important firstly because much research effort has been devoted to identifying metabolite markers of poor graft union formation and graft incompatibility with limited success (Loupit & Cookson, 2020), and secondly, because many genes related to secondary metabolite formation are induced during graft union formation (Assunção, Santos, Brazão, Eiras-Dias, & Fevereiro, 2019; Cookson et al., 2013) and the function of these metabolites is unknown.

### trans-Ε-viniferin is the major stilbene in grapevine canes and is heterogeneously distributed in grapevine stem tissues

The secondary metabolite found in highest concentration in the tissues studied was *trans*-ε-viniferin, a dehydrodimer, which is formed by the oxidative dimerization of resveratrol. *trans*-ε-viniferin was also found to be the major stilbene in other studies on the graft interface of grapevine (Loupit et al., 2022; Prodhomme et al., 2019), as well as in grapevine canes (Billet et al., 2018; Loupit et al., 2020). *trans*-ε-viniferin, like other stilbenes, is known to be a phytoalexin, and therefore to have strong antioxidant and anti-fungal capacities (Chong et al., 2009). The high levels of stilbenes present in the woody tissues of grapevine could function as a constitutive protection mechanism for these perennial tissues against both abiotic stresses and pathogens.

### Grapevine genotypes have different wood metabolite profiles

We have previously shown that there is a large degree of variation in cane metabolite profile across different *Vitis spp.* (Loupit et al., 2020). However, the three genotypes studied had broadly similar metabolite profiles except that 140Ru has lower stilbene concentrations than the other two genotypes studied, which in particular was due to its relatively lower concentration of stilbene dimers. This is because we studied grafting in commercially grafted scion and rootstock genotypes, which corresponds to a relatively limited range of genetic diversity. MALDI-MSI showed that stilbenes were most concentrated in the bark, pith and between the pith and xylem zones. In RGM and PN wood, stilbene dimers and α-viniferin were distributed throughout the pith, whereas in 140Ru these metabolites were largely restricted to the pith/xylem boundary.

In the wood of poplar, phenylalanine, one of the central amino acids involved in the phenylpropanoids biosynthesis, is localised in the cambium and new xylem formed, potentially explaining the high concentrations of stilbenes in the xylem part (Abreu et al., 2020). However, catechin, which come from phenylalanine pathway as well, was mainly localised in the phloem and cambium tissue of grapevine and largely absent from the xylem area, in agreement with results on poplar in which catechin was also absent from the xylem area (Abreu et al., 2020).

### Some metabolites changes over time, which is presumably related to spring reactivation of growth

The concentrations of the different compounds in the intact control cuttings were not significantly different between the parts above, below and at the interface, which indicates that the sampling zone under the bud has no influence on metabolite quantification. However, a slight increase in stilbenes and some other compounds was observed over time in the three areas sampled. An increase in stilbene concentration over time was also observed in the wood above and below the graft interface in the grafts of PN and RGM, whereas these metabolites did not change in the wood above and below for 140Ru homo-grafts and hetero-grafts. During the 15 DAG, there is a reactivation of the metabolism due to the breaking of dormancy and bud burst, which involves many changes in primary and secondary metabolism, and metabolism-related gene expression. For example, the transcript abundance of 12 stilbene synthases increases after the transfer of grapevine cuttings to warm dormancy release conditions (Noronha et al., 2021). The genotype-specific changes in cane metabolite concentration could be related to their responses to spring reactivation of growth; *V. vinifera* (PN) and *V. riparia* (RGM) are considered as low chill requiring, rapid bud bursting grapevine species, whereas *V. rupestris* (one of the parents of 140Ru) generally have higher chill requirements and slower bud break (Londo & Johnson, 2014).

### Grafting and wounding induce a rapid and transient accumulation of naringenin and taxifolin, which are mainly present in callus

The concentration of naringenin and taxifolin rapidly and transiently increased at the graft interface/wounding site, these compounds are intermediate metabolites and precursors in flavonoids biosynthesis. Naringenin and taxifolin are localised in the callus tissues 16 DAG, but as callus tissues are absent in the first few days after grafting, they presumably accumulate in the tissues damaged during wounding produce by grafting process. In addition to being intermediates in the synthesis of flavonoids, naringenin has also been identified as an allelochemical in annual species, as an activator of reactive oxygen species (ROS), salicylic acid and pathogen resistance, and as a signal molecule involved in root symbiosis with microorganisms (Dardanelli et al., 2010). The application of naringenin to roots inhibits vegetative growth and decreases lignin content (Deng, Aoki, & Yogo, 2004). Although naringenin treatment to a culture solution of bean seeds improved tolerance to oxidative stress by inducing some enzymes with antioxidant activity, such as superoxide dismutase (SOD) and catalase (CAT) decreasing ROS content and reducing lipid peroxidation (Ozfidan-Konakci, Yildiztugay, Alp, Kucukoduk, & Turkan, 2020; Yildiztugay, Ozfidan-Konakci, Kucukoduk, & Turkan, 2020). This could suggest that naringenin could play a role during the early stages of graft union formation by increasing oxidative stress tolerance after wounding or grafting. In Norway spruce, taxifolin is synthesized by a flavanone-3-hydroxylase enzyme and induced resistance to bark beetles (Hammerbacher et al., 2019). Furthermore, taxifolin accumulates in developing xylem and phloem around cambial zone (Hammerbacher et al., 2019), consistent with our results showing the presence of taxifolin in callus, where new vascular connexions are actively being formed. Recently, An *et al*. (2023) proposed a regulatory network regulating flavonoid accumulation under Postharvest Physiological Deterioration (PPD) in tuberous roots of *Manihot esculenta* Crantz, and observed that naringenin concentrations increased during PPD (An et al., 2023).

### Flavanols do not seem to play a role during the first 2 weeks after wounding/grafting

A significant decrease in flavanol concentration was observed after wounding cuttings and at the graft interface of the two hetero-grafts compared to surrounding tissues, while the sum of total flavanols was stable over time in the intact control. Lower concentrations of epicatechin at the graft interface in comparison to the surrounding woody tissues one month after grafting has been reported previously (Prodhomme et al., 2019). This could be because these metabolites are oxidised and degraded in response to wounding or grafting. It is therefore possible that a decrease in epicatechin content at the graft interface is due to its role as ROS-scavenging and as an antioxidant compound but is not synthesized fast enough to maintain tissue epicatechin concentrations. Some studies in grapevine (Assunção, Pinheiro, et al., 2019; Canas et al., 2015) and in pear or apricot (Hudina et al., 2014; Musacchi et al., 2000; Usenik et al., 2006) have indicated that catechin and epicatechin could be used as markers of graft incompatibility. For example, the study of Canas *et al*. (2015) suggested that catechin could be an incompatibility marker for incompatible Syrah clones one month after grafting (Canas et al., 2015), however we did not observe a spatial distribution of catechin at the graft interface consistent with this hypothesis. Furthermore, we did not identify significant differences in flavanol concentration or distribution between the two hetero-graft combinations studied, which are known to have different grafting success rates. The lack of agreement between these results could be because different types of graft failure (i.e. incompatibly related to the scion Syrah or poor grafting success with the rootstock 140Ru) could lead to the production of different metabolites at the graft interface.

### Gallate-compounds accumulate over the time and in the callus

Although most flavanols did not accumulate at the graft interface, flavanols with a gallate residue, such as epigallocatechin gallate, did accumulate at the graft interface and were shown to accumulate particular in the callus tissue. Flavanols are also known for their strong antioxidant capacity and it has already been shown that epigallocatechin, epigallocatechin gallate or even epicatechin gallate have higher antioxidant capacities than flavanol monomers or gallic acid alone (Rice-Evans, 1995). However, it was also interesting to note that gallic acid was found in higher content in callus and in developing tissues such as the cambium at 15 and 30 DAG compared to other wood parts (data not shown), indicating that it is also likely that a high concentration of flavanol gallates is related to a high content of gallic acid. No accumulation of flavanol gallates was reported on leaf after wound response during the first hours and days (Chitarrini, Zulini, Masuero, & Vrhovsek, 2017), while we were able to detect a rapid accumulation, even when no callus was formed. This could suggest that wounding responses in leaves and woody stems induces the production of different defence metabolites.

### Stilbene accumulation in wounded/grafted tissues is related to the size of the wound

The sum of total stilbenes increased in response to both wounding and grafting in all genotypes studied. Total stilbenes (as well as monomers and the different stilbene oligomers) accumulated to higher levels in the PN/PN homo-grafts than in the wounded PN cuttings relative to the intact cuttings, indicating that the larger the wound the higher the metabolite accumulation. This agrees with the observation that stilbenes accumulate along the graft interface in the tissues damaged by the grafting process, in particular the xylem parenchyma, so the larger the wound the more stilbenes will be accumulated. In perennial species, secondary metabolites are known to be accumulated in xylem parenchyma in response to abiotic or biotic stresses (Morris et al., 2020). This metabolic response is one part of the Compartmentalisation of Decay in Trees (CODIT) Model, which aims to limit pathogen infections in perennial plant structures. The CODIT model considers that the xylem parenchyma is an important barrier to pathogens and produces phytoalexins in response to stress, limiting or compartmenting ROS and pathogen infections (Słupianek, Dolzblasz, & Sokołowska, 2021). The accumulation of defence related compounds after wounding or grafting could have a role in protecting the healthy tissues from pathogen infection. The wood metabolite profile of grapevine, in particular the ability to produce resveratrol and viniferins after wounding, has been linked to resistance to Botryosphaeriaceae-related dieback (Khattab et al., 2021), suggesting that these metabolite differences have consequences for plant function.

### Stilbenes accumulate after wounding/grafting and oligomerize over time

At the end of the time course, stilbene monomer concentrations decreased at the graft interface (from 3 or 6 to 14 DAG); this could suggest that these monomers were used to form oligomers over time. The sequential appearance of stilbene monomers and then oligomers was also found in a study on metabolite changes in grapevine leaves after wounding (Chitarrini et al., 2017). The biosynthetic pathway of complex compounds is currently unknown, which does not allow us to know whether these compounds are actively formed by the plant, or passively accumulated. One of the hypotheses would be that the monomers oligomerize over time via their oxidation. The synthesis of resveratrol dimer has been carried out using *in vitro* oxidative coupling (El Khawand et al., 2020). In addition, oxidative oligomerization can occur via enzymatic reactions with horseradish peroxidase (Wilkens, Paulsen, Wray, & Winterhalter, 2010) or laccase-like stilbene oxidase from Botrytis cinerea (Breuil et al., 1998), although there is no direct evidence of these enzymes catalysing the oxidative oligomerization of stilbenes in plants.

### Metabolite markers of graft compatibility

Few differences were found in stilbene concentration between the two hetero-grafts with very different grafting success rates (60.3% for PN/RGM and 20.3% for PN/140Ru), except that higher pallidol and miyabenol C concentrations were measured in PN/RGM compared to PN/140Ru. A similar trend was also observed in homo-grafts with higher grafting success rates (RGM/RGM and 140Ru/140Ru had grafting success rates of 28.1 and 32.8 % respectively) in comparison with PN homo-grafts with lower grafting success rates (6.25 %). Conversely, α-viniferin was found in higher concentrations in grafts with low grafting success rates. In agreement with this observation, we have shown that this trimer is negatively correlated with grafting success in homo-graft combinations and it was identified as a potential graft incompatibility marker (Loupit et al., 2022). In addition, resveratrol concentration at the graft interface one month after grafting was also previously identified as a compound positively correlated with grafting success (Loupit et al., 2022). We also found that resveratrol increased to higher concentrations in the hetero-graft with this highest grafting success (PN/RGM) compared to the hetero-graft with lower grafting success (PN/140Ru) at 3-6 DAG. However, the concentration of resveratrol 15 DAG was not different between the different scion/rootstock combinations. Identifying robust and stable metabolite markers of grafting success or compatibility is challenging (Loupit & Cookson, 2020), the spatiotemporal characterisation of metabolite changes occurring during graft union formation underlines the complexity of this challenge.

## Conclusion

The objective of this study was to resolve the temporal changes in secondary metabolite concentration underlying graft union formation and to characterise the spatial distribution of these metabolites. Few studies have looked at the kinetics of the accumulation of secondary metabolites in response to wounding/grafting in grapevine except for one study on metabolites changes in leaf discs during the first 120 hours after wounding (Chitarrini et al., 2017). The comparison of metabolites accumulated after wounding in leaf and woody stem tissues identified some metabolites specific to woody stem wounding responses such as naringenin, taxifolin and flavanol gallate accumulation. In this study we have shown that stilbenes are highly and rapidly accumulated at the graft interface and in response to wounding, and that these stilbenes oligomerize over time. We have also shown that the many other metabolites show large temporal changes in concentration at the graft interface, particularly naringenin and taxifolin. In addition, knowledge of the spatial distribution of metabolites accumulated after wounding or grafting provides insights into their potential roles (summarized in Figure 7). For example, metabolites which accumulate in the damaged xylem parenchyma such as stilbenes have potential roles in plant defence to prevent pathogen infection into perennial woody structures. Whereas secondary metabolites which accumulate in the newly formed callus tissues presumably have a wider range of roles such as forming new vascular tissues, plant signalling, plant defence and developing a functional graft union.

**Figure 7:**
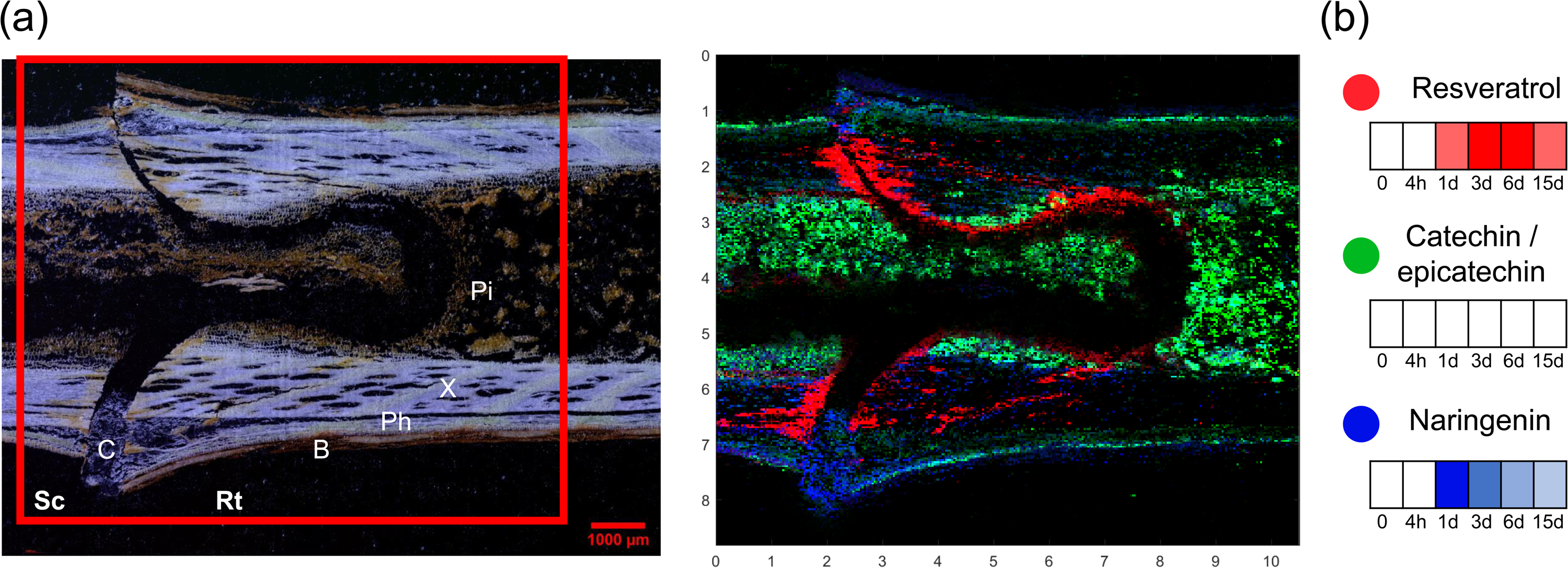
Summary of the (a) spatial and (b) temporal changes in resveratrol, catechin/epicatechin and naringenin at the graft interface during the first two weeks after grafting. (a) The section shown is of *V. vinifera* cv. Pinot Noir/*V. berlandieri x V. rupestris* cv. 140 Ruggeri., MS image generated for *m/z* 227.0708, *m/z* 289.0712 and *m/z* 271.0607 corresponding to resveratrol (red), catechin/epicatechin (green) and naringenin (blue) respectively. The red box corresponds to the area analysed. Sc, scion; Rt, rootstock; C, callus; B, bark; Ph, phloem; X, xylem; Pi, pith. (b) Heatmaps of metabolite concentration at the graft interface relative to the surrounding woody tissues during the first 2 weeks after grafting.

## Acknowledgements

Research support was provided by the French Ministry of Higher Education, and by the European Union INTERREG POCTEFA project Vites Qualitas (EFA 324/19) which is co-financed by the Fonds Européen de Développement Regional (FEDER). Also, this article is based upon work from COST Action CA17111 INTEGRAPE, supported by COST (European Cooperation in Science and Technology). This work was supported by the Bordeaux Metabolome Facility (https://doi.org/10.15454/1.5572412770331912E12) and the MetaboHUB (ANR-11-INBS-0010) project. Support from the Carlsberg Foundation and The Danish Council for Independent Research|Medical Sciences (Grant DFF-4002-00391) for the applied MALDI-MSI instrumentation is gratefully acknowledged.

We thank Cyril Hevin, Maria Lafargue and Nicolas Hocquard for the grafting plants and Anne Janoueix, Marilou Camboué, Pablo Dupiol and Fernanda Endringer Pinto for helping during the sampling and sample preparation.

## Author contribution

SJC, JVF and GL design the experiments; GL performed sample preparations and extractions. JVF, CF, GdeR and GL performed secondary metabolites analysis by HPLC-QqQ analysis; ML, CJ and GL performed MALDI-MSI analysis; GL analysed, interpreted and performed statistical analysis of the data; GL wrote the manuscript; JVF and SJC revised and corrected the manuscript. All authors read and approved the final manuscript.

## Data availability

All quantification data (means and standard deviation) all conditions studied are given in Supplementary data 1. MALDI-MSI data is available on METASPACE annotations (https://metaspace2020.eu/project/Loupit-2023) (Palmer et al., 2017).

## Supporting Information

Data S1: Effect of grafting and wounding on stilbenes, flavanols, flavonols and phenolic acids at 0 and 4 h, and 1, 3, 6 and 15 d after grafting/wounding, above, below and at the graft interface. Cuttings and grafts as described in legend for Fig. 1. Means shown, errors bars represent the standard deviation (n=5 for grafts and n=3 for cuttings).

Data S2: *p*-Values found for differences in metabolite concentration between above and at the graft interface, and between below and at the graft interface in the cuttings and grafts described in legend for Fig. 1., harvested at 0 and 4 h, and 1, 3, 6 and 15 d after grafting/wounding. Excel cells in red show *p*-value < 0.05.

Table S1: Mass of precursor ion (Precursor Ion), mass of one of the fragments used for quantification (Quantifier), retention time (Rt) and ion polarity for the 41 secondary metabolites for HPLC-QqQ in MRM mode.

Table S2: MALDI images (first biological replicate) of V. berlandieri x V. rupestris cv. 140 Ruggeri wood at 0 and homo-grafts at 16 and 30 d after grafting. Images show the distribution of resveratrol, stilbene dimers, α-viniferin, stilbene tetramers, flavanol monomers (catechin/epicatechin), flavanol dimers, epigallocatechin gallate, taxifolin and naringenin compounds observed at m/z 227.0708, 453.1338, 667.1812, 905.2598, 289.0712, 577.1346, 457.0771, 303.0505 and 271.0607 respectively. The red boxes correspond to the area analysed.

Table S3: MALDI images (second biological replicate) of V. berlandieri x V. rupestris cv. 140 Ruggeri wood at 0 and homo-grafts at 16 and 30 d after grafting. Images as described in Table S2.

Table S4: MALDI images (first biological replicate) of *V. vinifera* cv. Pinot Noir wood at 0 and homo-grafts at 16 and 30 d after grafting. Images show the distribution of resveratrol, stilbene dimers, α-viniferin, stilbene tetramers, flavanol monomers (catechin/epicatechin), flavanol dimers, epigallocatechin gallate, taxifolin and naringenin compounds observed at *m/z* 227.0708, 453.1338, 667.1812, 905.2598, 289.0712, 577.1346, 457.0771, 303.0505 and 271.0607 respectively. The red boxes correspond to the area analysed.

Table S5: MALDI images (second biological replicate) of V. vinifera cv. Pinot Noir wood at 0 and homo-grafts at 16 and 30 d after grafting. Images as described in Table S4.

Table S6: MALDI images (first biological replicate) of *V. riparia Michaux* cv Riparia Gloire de Montpellier wood at 0 and homo-grafts at 16 and 30 d after grafting. Images show the distribution of resveratrol, stilbene dimers, α-viniferin, stilbene tetramers, flavanol monomers (catechin/epicatechin), flavanol dimers, epigallocatechin gallate, taxifolin and naringenin compounds observed at *m/z* 227.0708, 453.1338, 667.1812, 905.2598, 289.0712, 577.1346, 457.0771, 303.0505 and 271.0607 respectively. The red boxes correspond to the area analysed.

Table S7: MALDI images (first biological replicate) of V. riparia Michaux cv Riparia Gloire de Montpellier wood at 0 and homo-grafts at 16 and 30 d after grafting. Images as described in Table S6.

Table S8: MALDI images (first biological replicate) of *V. vinifera* cv. Pinot Noir / *V. berlandieri x V. rupestris* cv. 140 Ruggeri hetero-grafts at 16 and 30 d after grafting. Images show the distribution of resveratrol, stilbene dimers, α-viniferin, stilbene tetramers, flavanol monomers (catechin/epicatechin), flavanol dimers, epigallocatechin gallate, taxifolin and naringenin compounds observed at *m/z* 227.0708, 453.1338, 667.1812, 905.2598, 289.0712, 577.1346, 457.0771, 303.0505 and 271.0607 respectively. The red boxes correspond to the area analysed.

Table S9: MALDI images (first biological replicate) of V. vinifera cv. Pinot Noir / V. berlandieri x V. rupestris cv. 140 Ruggeri hetero-grafts at 16 and 30 d after grafting. Images as described in Table S8.

Table S10: MALDI images (first biological replicate) of *V. vinifera* cv. Pinot Noir / *V. riparia Michaux* cv Riparia Gloire de Montpellier hetero-grafts at 16 and 30 d after grafting. Images show the distribution of resveratrol, stilbene dimers, α-viniferin, stilbene tetramers, flavanol monomers (catechin/epicatechin), flavanol dimers, epigallocatechin gallate, taxifolin and naringenin compounds observed at *m/z* 227.0708, 453.1338, 667.1812, 905.2598, 289.0712, 577.1346, 457.0771, 303.0505 and 271.0607 respectively. The red boxes correspond to the area analysed.

Table S11: MALDI images (first biological replicate) of V. vinifera cv. Pinot Noir / V. riparia Michaux cv Riparia Gloire de Montpellier hetero-grafts at 16 and 30 d after grafting. Images as described in Table S10.

Table S12: MALDI images (first biological replicate) of *V. riparia Michaux* cv Riparia Gloire de Montpellier wood at 16 and 30 d after grafting. Images show the distribution of resveratrol, stilbene dimers, α-viniferin, stilbene tetramers, flavanol monomers (catechin/epicatechin), flavanol dimers, epigallocatechin gallate, taxifolin and naringenin compounds observed at *m/z* 227.0708, 453.1338, 667.1812, 905.2598, 289.0712, 577.1346, 457.0771, 303.0505 and 271.0607 respectively. The red boxes correspond to the area analysed.

Table S13: MALDI images (first biological replicate) of V. riparia Michaux cv Riparia Gloire de Montpellier wood at 16 and 30 d after grafting. Images as described in Table S12.

